# Multi-disease associated risk locus in *IL6* represses the anti-inflammatory gene *GPNMB* through chromatin looping and recruiting MEF2-HDAC complex

**DOI:** 10.1101/602797

**Authors:** Xiufang Kong, Amr H Sawalha

## Abstract

We have previously revealed a genetic association between Takayasu arteritis and a non-coding genetic variant in an enhancer region within *IL6* (rs2069837 A/G). The risk allele in this variant (allele A) has a protective effect against chronic viral infection and cancer. Using a combination of experimental and bioinformatics tools, we identified the monocyte/macrophage anti-inflammatory gene *GPNMB*, ∼520kb away, as a target gene regulated by rs2069837. We revealed preferential recruitment of myocyte enhancer factor 2-histone deacetylase (MEF2-HDAC) repressive complex to the Takayasu arteritis risk allele. Further, we demonstrated suppression of GPNMB expression in monocyte-derived macrophages from healthy individuals with the AA compared to AG genotype, which was reversed by histone deacetylase inhibition. Our data suggest that the A allele in rs2069837 represses the expression of GPNMB by recruiting MEF2-HDAC complex, enabled through a long-range intra-chromatin looping mediated by CTCF. Suppression of this anti-inflammatory gene might mediate increased susceptibility in Takayasu arteritis and enhance protective immune responses in chronic infection and cancer. Our data highlight long-range chromatin interactions in functional genomic studies.

## Introduction

Takayasu arteritis is a granulomatous large vessel vasculitis, mainly affecting women of childbearing age (*1*). The disease primarily involves the aorta and its major branches, leading to thickening, stenosis, and occlusion of involved vessels. Takayasu arteritis is relatively more prevalent in Asia and North Africa compared with Europe and North America (*2*).

Although Takayasu arteritis occurs worldwide, the etiology of this disease remains elusive, in part due to its rarity and indolent course. It is suggested that both genetic and environmental factors contribute to the development of Takayasu arteritis (*2*). Several genetic susceptibility loci have been identified and confirmed in Takayasu arteritis, however, there is no convincing evidence regarding the specific environmental factors that could be involved. The genetic association between Takayasu arteritis and *HLA-B*52* has been confirmed in multiples cohorts and ethnicities (*3*). In addition, non-HLA susceptibility loci including *FCGR2A*/*FCGR3A, IL12B, IL6, RPS9*/*LILRB3*, have been reported with a genome-wide level of significance (*4-6*).

Similar to other autoimmune diseases, the majority of genetic susceptibility loci reported in Takayasu arteritis in genome-wide association studies are in non-coding regions. In addition, causal genetic variants in these loci and the target genes affected are largely unknown. Given the complex and three dimensional nature of the human genome, genetic susceptibility loci do not necessarily affect genes within which they are located or to which they are closest, but instead can potentially affect other target genes at a distance. Therefore, identifying causal variants and uncovering the potential regulatory effects within these genetic susceptibility loci and affected cell types are critical to revealing target genes and elucidating the mechanisms upon disease susceptibility.

We have previously identified a genetic association between *IL6* and Takayasu arteritis (*4*). Specifically, we reported a genetic association between a locus tagged by rs2069837(A/G) located within the second intron of the *IL6* genes, with allele A being the disease risk allele in Takayasu arteritis. This same genetic variant has been reported to be associated with multiple other diseases and conditions. The Takayasu arteritis associated risk allele in this variant is associated with longevity, spastic tetraplegia, cerebral palsy, and antipsychotic-induced weight gain (*7-10*). In contrast, the Takayasu arteritis risk allele in rs2069837 was shown to be protective against late-onset Alzheimer’s disease, cervical cancer, chronic hepatitis B virus infection, colorectal cancer, and hepatocellular carcinoma (*11-15*).

In the present study, we characterize the regulatory function of rs2069837 and reveal a distant target gene affected by this genetic polymorphism. The findings of this study mechanistically elucidate the genetic effect of this variant relevant to multiple disorders.

## Results

We have previously revealed a genetic association between Takayasu arteritis and rs2069837, which is located in the second intron of *IL6* (hg19, chr7: 22768026). No genetic variants in linkage disequilibrium with rs2069837 (r^2^>=0.6) were identified, suggesting that rs2069837 is the likely causal variant in this locus (*4*). DNase hypersensitivity and acetylation of histone H3 on lysine 27 (H3K27ac) data in various immune cells suggest that this genetic variant is near an enhancer region (**Figure 1A**). To identify regulatory proteins that bind rs2069837 we first performed an electrophoretic mobility shift assay (EMSA) using DNA oligos with the Takayasu arteritis risk allele (allele A) and the protective allele (allele G) in rs2069837. We show that more DNA-protein complexes were formed in the presence of probe with allele A than allele G (**Figure 1B**), which was also confirmed using nuclear extract from THP-1 cells (**Supplementary Figure 1A**). Both complexes can be competed out by their unlabeled probes respectively. In addition, unlabeled probe with A allele was able to compete the shift band formed in the presence of probe with G allele in both cell lines (**Figure 1B, Supplementary Figure 1A**). Further, when we used different folds of unlabeled probe with allele A to compete labeled probe with allele A, a 10-fold concentration (1.2 pmol) of unlabeled probe with allele A was sufficient to completely compete with the shift band, suggesting that the binding affinity between DNA sequence and nuclear proteins was high (**Figure 1C**).

**Figure 1.**
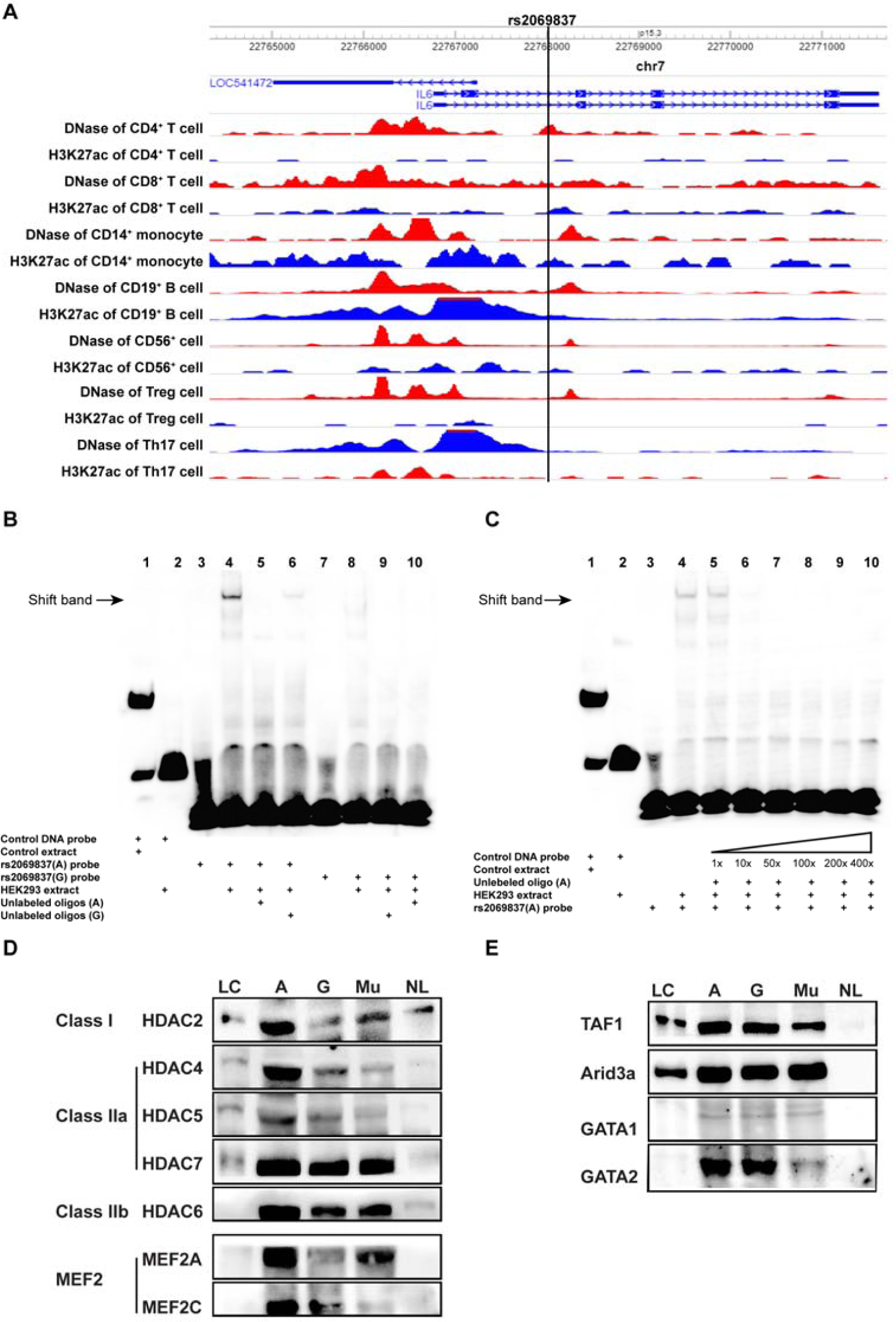
Characterization of rs2069837 and detection of binding proteins. A. rs2069837 (hg19, chr7: 22768026) is located in the second intron of *IL6*. The epigenetic markers (DNase and H3K27ac) in multiple immune cells indicate that this locus is close to a functional region. B. EMSA assay demonstrating that more DNA-protein complexes were shifted with the probe with allele A (lane 4) than allele G (lane 8); both unlabeled probes with A allele or G allele (24 pmol, 200X) can totally compete the corresponding shift band separately (lane 5 and lane 9); unlabeled probe with G allele cannot compete the shifted band formed with the probe with A allele (lane 6); unlabeled probe with A allele was able to compete the shifted band formed in the presence of probe with G allele (lane 10). C. Different folds of the unlabeled probe with A allele were applied in the competitive EMSA assay (1X, 10X, 20X, 50X, 100X, 200X). At over 10X concentration, the band can be competed out completely (lane 6). D-E. Validation using Western blotting. More HDAC and MFE2 proteins bound to the probe with A than with G allele, while there was no differences were detected in TAF1, ARID3A, GATA1, and GATA2 between probes with A and G. When the four nucleotides around rs2069837 were muted, the binding of those proteins decreased except for ARID3A. The protein binding in the unlabeled group was barely observed (LC, loading control; A: probe with A allele; G: probe with G allele; mu: probe with muted sequence; NL: non-biotinylated oligos with A allele).

To investigate possible regulatory proteins that bind rs2069837, we first examined publically available data using HaploReg and CIS-BP databases (*16, 17*). Publically available experimental data indicate that rs2069837 can alter motifs of several important transcription regulators (**Supplementary Table 3**). The changes of log-odds (LOD) scores between reference allele A and alternative allele G indicated alterations of binding affinities for corresponding transcription regulators. The scores were increased for all the binding motifs, except TATA_known3, from G allele to A allele at this SNP, indicating that the risk allele A potentially binds more proteins, consistent with our EMSA results. Among them, the binding affinities of MEF2 and FOXJ1 increased the most, followed by ARID3A, FOXA, FOXD3, FOXF1, and FOXI1. When we used CIS-BP database to predict potential binding proteins, we identified ARID3 proteins, Forkhead family proteins, MEF2 and GATA proteins. Among these, ARID3 and MEF2 proteins were shown only in the sequence with A allele, while GATA proteins were predicted to bind only the sequence with G allele (**Supplementary Table 4**).

Next, we used DNA affinity precipitation followed by mass spectrometry to confirm proteins that differentially bind to the motif around rs2069837 in the presence of A and G alleles **(Supplementary Table 5)**. MEF2 proteins were only detected in proteins bound to DNA with A allele, while more GATA proteins appeared in complexes with G allele, which is consistent with bioinformatics prediction and mining publically available data. However, no remarkable differences were observed in the binding of ARID family, forkhead family, and TATA-binding proteins associated factors between A and G alleles. In addition, histone deacetylases (HDACs) were also detected in these protein complexes, among which HDAC5 was mainly detected in complex with allele A. These results were further validated by Western Blotting which showed increased MEF2 and HDAC binding in the presence of the A allele compared to the G allele, consistent with mass spectrometry results (**Figure 1 D and E**). It has been known that HDAC class IIa family such as HDAC4, HDAC5, HDAC7 have a unique regulatory domain which can mediate an interaction with MEF2 factors, causing the repression of MEF2-target genes. Taken together, our data indicate that the Takayasu arteritis risk allele (allele A) in rs2069837 induces transcriptional repression due to preferential recruitment of the MEF2-HDAC complex, thereby weakening the enhancer function in this locus.

To confirm that rs2069837 is in an enhancer region and that allele A is associated with transcriptional repression, we utilized a luciferase reporter assay (**Figure 2A**). After normalizing to renilla luciferase activity, both plasmids inserted with the intronic sequence containing A or G showed significantly higher luciferase activity compared with the basic vector or promoter-only vector (P < 0.05, **Figure 2B**). When comparing the luciferase activity between the plasmid with the insert containing A allele and the plasmid with the insert containing G allele, the plasmid with G allele demonstrated stronger luciferase activity (P <0.05, **Figure 2B**). These data were also confirmed in THP-1 cell line (**Supplementary Figure 1B**), demonstrating that rs2069837 is in an enhancer region, which is significantly weakened in the presence of the Takayasu arteritis risk allele (allele A), consistent with our findings above. To test the hypothesis that allele A induces transcriptional repression by preferentially recruiting MEF2-HDAC complex, we assessed the effect of HDAC inhibitor TSA on luciferase activity in the presence of the A and G alleles. With TSA treatment, the luciferase activity was increased in both constructs with A and G alleles, confirming recruitment of HDAC to rs2069837. Importantly, luciferase activity was significantly more increased with TSA treatment in the presence of the A allele, confirming a functional consequence of the preferential recruitment of MEF2-HDAC complex to allele A in rs2069837 (**Figure 2C**).

**Figure 2.**
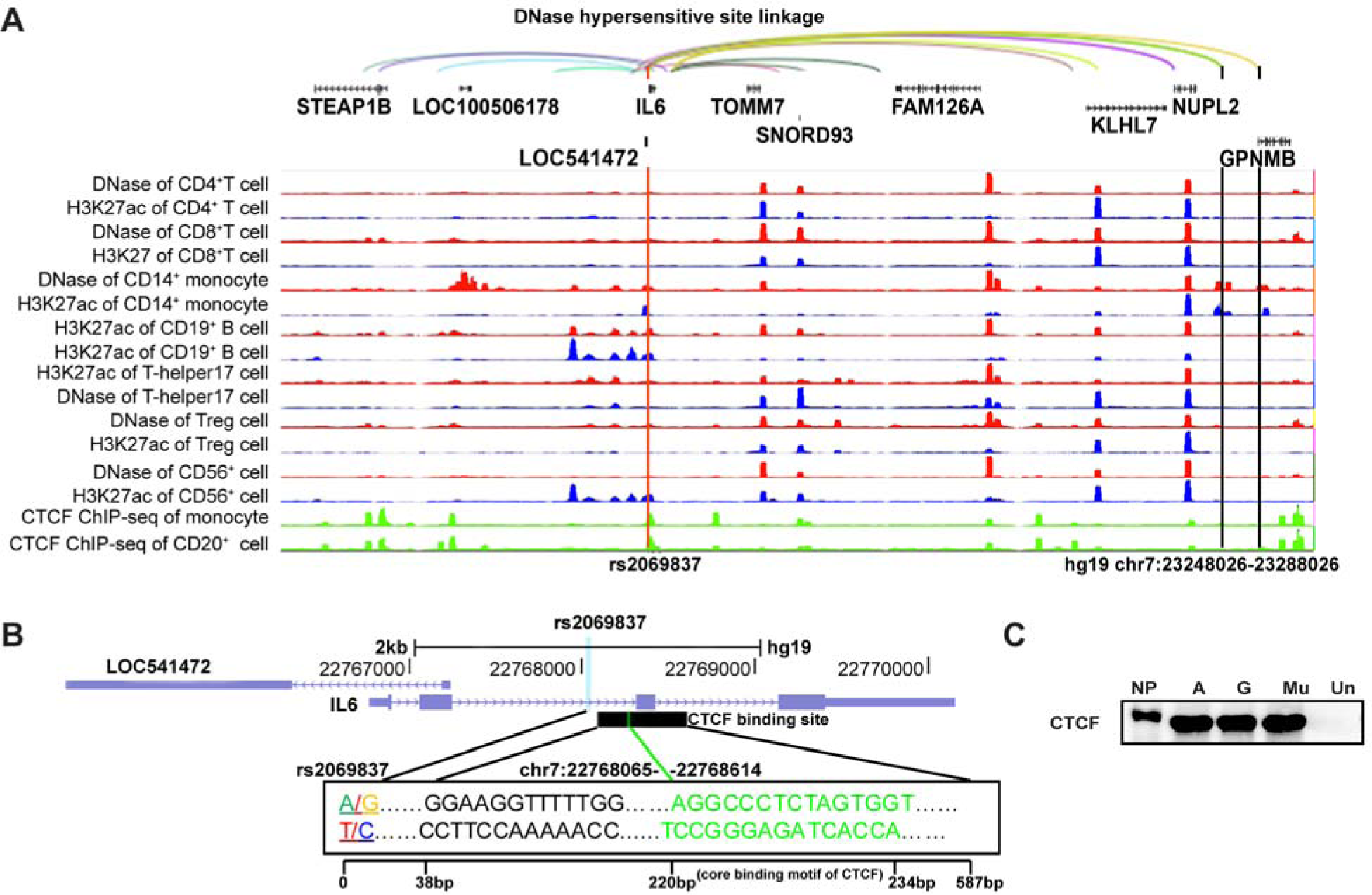
Luciferase reporter assay to assess enhancer function in rs2069837 and the effect of inhibiting HDAC. A. The intronic sequence with A or G (hg19 chr7:22767927-22768177) was inserted between Xba I and EcoRI, while the promoter sequence (nucleotides–303 to +12, Ensembl ENSG00000136244) was inserted between Spe I and Mlu I. B. No increase of luciferase expression was observed in promoter only vector. Significantly increased luciferase expression was detected in both vectors with intronic sequence (A/G), with significantly higher expression in the construct with G allele than that with A allele (p< 0.05). C. TSA treatment increased luciferase expression in both constructs with intronic sequence (A/G) compared to DMSO treated samples. In the presence of TSA, the luciferase expression was significantly higher in the construct with A allele than that with G allele (p< 0.05).

To identify target genes regulated by the rs2069837 locus, we examined chromatin interactions involving rs2069837 using DNAase hypersensitivity site (DHS) linkage patterns across various tissues (*18, 19*). Multiple DHS linkages were observed between this SNP and distal loci DHS (hg19, chr7: 22528026, chr7:22568026, chr7:22688026, chr7:22968026, chr7:23128026, chr7:23228026, chr7:23248-026, chr7:23288026). These interacting regions covered several genes, including *STEAP1B, LOC100506178, LOC541472, IL6, TOMM7, SNORD93, FAM126A, KLHL7, NUPL2, GPNMB* (**Figure 3A**). Since CTCF-mediated chromatin looping is the most common mechanism for long-distance chromatin interaction (*20*), we also checked the CTCF enrichment at this region using ChIP-seq data through WashU EpiGenome browser. ChIP-seq data for CTCF were only available in monocytes and B cells, and the rs2069837 locus was enriched for CTCF binding in both cell types (**Figure 3B**). Binding of CTCF to this locus was further validated by Western blotting and was not different between alleles A and G in rs2069837, and the mutated sequence, indicating that the binding of CTCF occurs in this locus but is not affected by rs2069837. Moreover, CTCF and units of cohesion complex, which is composed of multiple proteins that play important role in CTCF-mediated looping, were also detected in our mass spectrometry results (**Supplementary Table 6**).

**Figure 3.**
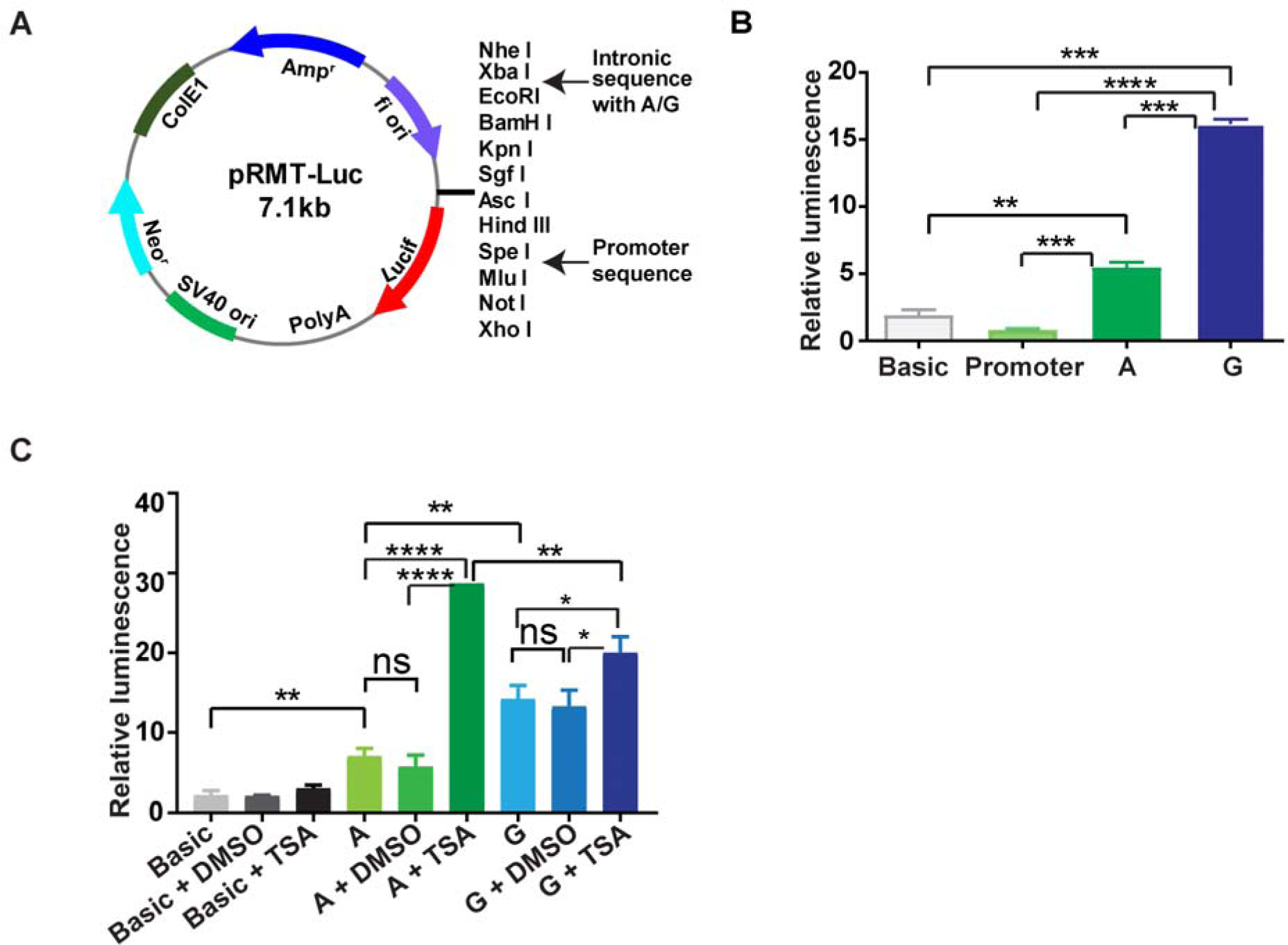
Potential interactions between rs2060837 and distant loci. A. DNase hypersensitivity linkage (40kb resolution) showed correlation of DNase hypersensitivities between rs2069837 and several distant loci. The interaction region (890kb) included several genes such as *STEAP1B, LOC100506178, LOC541472, IL6, TOMM7, SNORD93, FAM126A, KLHL7, NUPL2 and GPNMB* from 5’ to 3’. The interaction sites from upstream to downstream were chr7: 22528026, chr7:22568026, chr7:22688026, chr7:22968026, chr7:23128026, chr7:23228026, chr7:23248-026, chr7:23288026 (hg19). Among them, the last two interaction sites (chr7: 23248026, 23288026) overlapped with Dnase hypersensitivity and H3K27ac peaks in monocyte, where the closest gene was *GPNMB*. In addition, there is CTCF binding enrichment at rs2069837 in monocytes and B cells. B. CTCF binding sites downstream of rs2069837 (position: chr7:22768065-22768614, 38 to 587bp downstream of rs2069837; core binding motif: chr7:22768247-22768261, 220bp to 234bp downstream of rs2069837). The black bar/track shows regions of transcription factor binding derived from a large collection of ChIP-seq experiments through across multiple cell lines presented using genome browser. C. Western blot analysis demonstrating CTCF binding to the probes with A or G alleles. No difference between A, G, or muted sequence was detected, indicating the binding of CTCF was not affected by rs2069837 (LC: loading control; A: probe with A allele; G: probe with G allele; mu: probe with muted sequence; NL: non-biotinylated oligos with A allele).

Next, we explored whether the rs2069837 locus regulates *IL6* or other genes indicated by chromatin interaction loops using DHS data (*STEAP1B, LOC100506178, LOC541472, TOMM7, SNORD93, FAM126A, KLHL7, NUPL2, GPNMB*). First, we performed functional annotation analysis of these genes using DAVID. The results demonstrated that GPNMB has a close relationship with IL-6 and participates in multiple biological processes such as angiogenesis (**Supplementary Table 7**). No relevant significant finding was found for the other genes. By further literature review, we found that GPNMB is expressed by multiple cell types including monocytes and it is increasingly expressed when monocytes differentiate into macrophages. Because monocytes/macrophages play an important role in the pathogenesis of Takayasu arteritis (*21-24*), and that epigenetic patterns (DNase hypersensitivity and H3K27ac; **Figure 2A**) also indicated that the rs2069837-interacting site in the *GPNMB* locus is a potential regulatory region in monocytes rather than other immune cells, we used primary monocyte-derived macrophages for our subsequent functional studies.

Forty-eight normal healthy individuals were genotyped for rs2069837 (**Supplementary Figure 1C**). Among them, 7 were AG and the remaining subjects were AA genotypes. Thus, we selected 7 pairs of age and ethnicity-matched individuals with AG and AA genotypes for further studies (**Supplementary Table 8**). In monocyte-derived macrophages, we observed no differences in IL6 mRNA expression, while GPNMB expression was significantly lower in macrophages derived from subjects with AA compared to AG genotype (**Figure 4A**). No differential expression was observed in all other interacting genes between samples with AA and AG genotypes (**Supplementary Figure 1D**). Importantly, TSA treatment significantly enhanced GPNMB expression in individuals with AA genotype but not AG genotype (**Figure 4 B, C**). This is consistent with our data derived from the luciferase assays and confirms a role for preferential recruitment of MEF2-HDAC complex in regulating GPNMB expression in the presence of the A allele in rs2069837.

**Figure 4.**
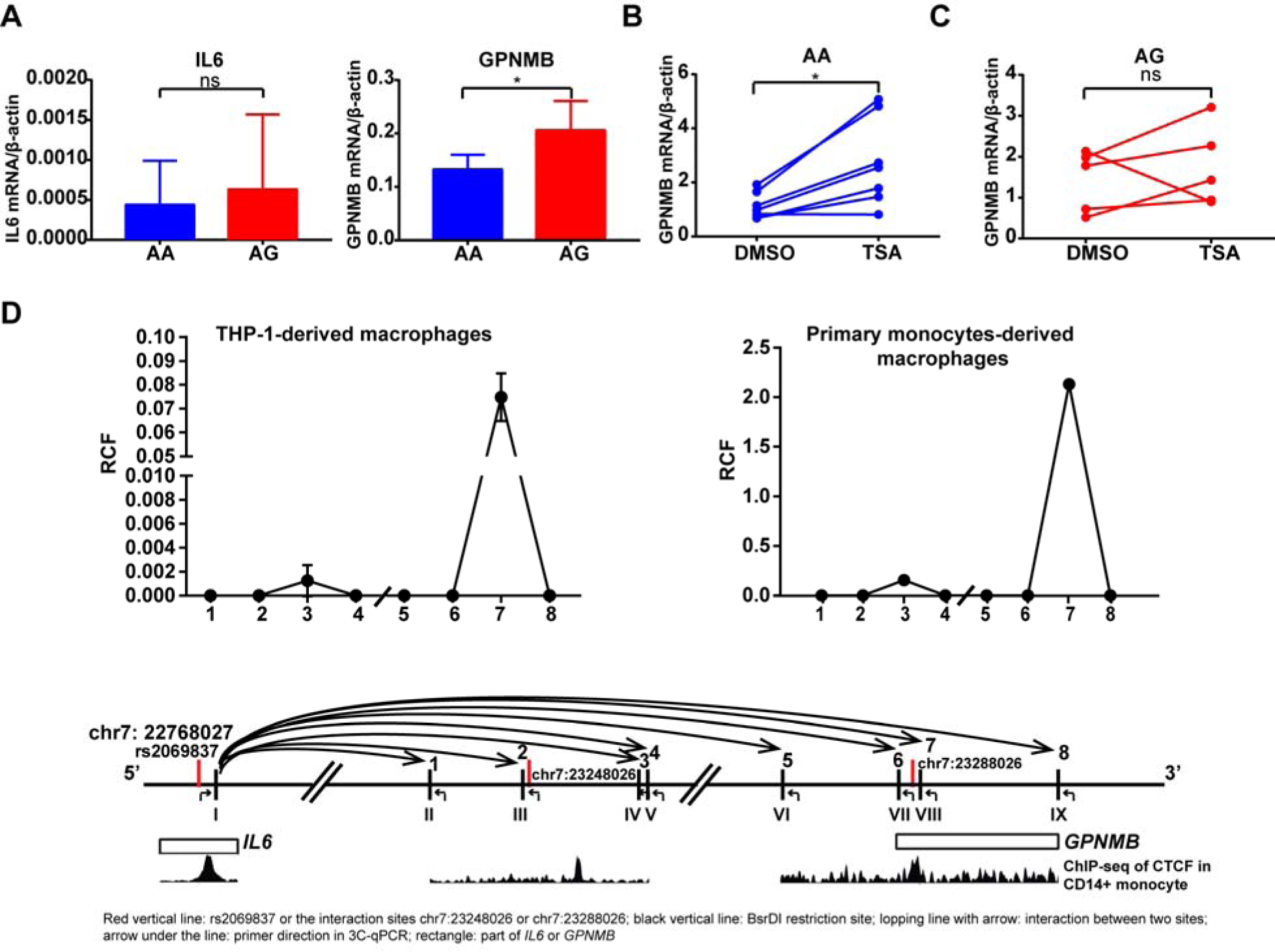
Detection of GPNMB as target gene repressed by the Takayasu arteritis risk allele in rs2069837 through long-distance chromatin looping. A. The mRNA expression level of GPNMB was significantly lower in monocyte-derived macrophages with AA compared to AG genotypes (p<0.05, n=7 in each group). No difference was observed in IL6 expression. B. TSA treatment (100nM) significantly increased GPNMB expression in monocyte-derived macrophages with AA genotype (p<0.05, n=7). C. TSA treatment (100nM) did not affect GPNMB expression in monocyte-derived macrophages with AG genotypes (p>0.05, n=5). D. 3C data revealed the interaction between rs2069837 and chr7:23288026 (*GPNMB*) in THP-1 cells (two replicates) and primary monocytes-derived macrophages (representative of two independent samples). The interactions examined relative to chromosomal positions are shown in the bottom panel. Restriction sites are depicted using Latin numbers, and interaction loops tested are depicted in Arabic numbers.ChIP-seq data of CTCF in CD14+ monocyte demonstrated that CTFC was strongly enriched at rs2069837 and chr7:23288026 (hg19) but weakly enriched at chr7:23248026 (hg19). The relative positions of restriction sites to chr7:23248026 or chr7:23288026 are as follows: I: +871bp of rs2069837 and +284bp of CTCF binding region, II: -7797bp of chr7:23248026, III: -29bp of chr7:23248026, IV: +6906bp of chr7:23248026, V: +7179bp of chr7:23248026, VI: -9293bp of chr7: 23288026, VII: -1933bp of chr7:23288026; VIII: +578bp of chr7:23288026; IX: +9224bp of chr7:23288026.

Further examination of the interaction between rs2069837 and *GPNMB* based on DHS linkage indicates two interactions, one 38.29kb upstream of *GPNMB* (hg19, chr7:23248026), while the other (hg19, chr7:23288026) was located in the first intron of *GPNMB* (**Figure 3A**). Both loci are within DNase and H3K27ac peaks in monocytes rather than other immune cells, indicating the regulatory function of these two loci in monocytes. CTCF ChIP-seq data also indicated peaks at these loci (**Figure 3A**). Other positive epigenetic marks (H3K4me1 and H3K4me1) in monocytes further demonstrated that these two rs2069837-interacting loci, especially chr7:23288026, are within important regulatory regions in *GPNMB* (**Supplementary Figure 1E**). In addition, ChIP-seq data in several cell lines showed GATA and MEF2 enrichment at these loci which were also demonstrated by analyzing the sequence motifs (**Supplementary Figure 1E**).

To confirm the interaction between rs2069837 and *GPNMB* (chr7:23288026), a 3C experiment was performed which demonstrated an interaction between the rs2069837 locus and a locus close to chr7:23288026 (**Figure 4D**). Since the BsrDI restriction sites close to rs2069837 and chr7: 23288026 were also close to the regions enriched with CTCF, CTCF was assumed to mediate this interaction.

Taken together, our data indicate that the Takayasu arteritis risk allele in rs2069837 represses an enhancer function in this locus by recruiting MEF2-HDAC complex and inhibits GPNMB expression via long-distance intra-chromatin looping likely mediated by CTCF (**Figure 5**).

**Figure 5.**
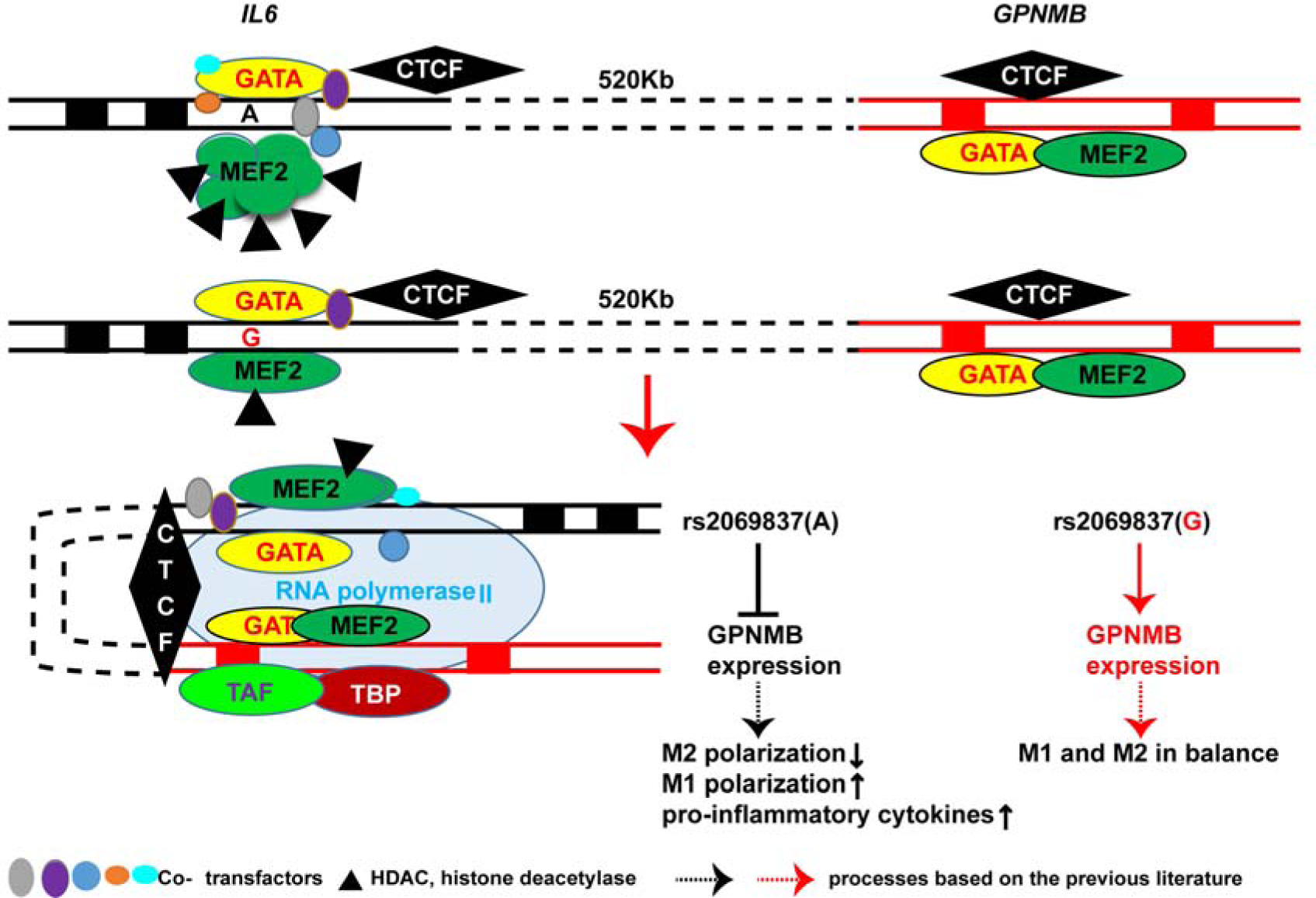
A proposed model for the regulatory mechanism of rs2069837 on the expression of *GPNMB*. The Takayasu arteritis risk allele A at this SNP preferentially recruits MEF2 and thereby HDAC proteins compared to the G allele resulting in a repressive effect by weakening the enhancer function of this locus. CTCF binding downstream of this SNP mediates the interaction between this locus to the regulatory locus in *GPNMB*, where a CTCF binding site also exists. The differential binding of MEF2-HADAC complex between A and G results in differential expression of GPNMB, with inhibited expression in the presence of risk allele A. Inhibiting GPNMB suppresses M2 macrophage polarization and enhances of M1 polarization and overexpression of pro-inflammatory cytokines.

## Discussion

In this study, we demonstrate that rs2069837 is in an enhancer region that is preferentially suppressed in the presence of the Takayasu arteritis risk allele, and although located within *IL6*, it affects the expression of a distant anti-inflammatory gene in human monocyte-derived macrophages. We show that the risk allele A recruits a repressive MEF2-HDAC protein complex, weakening the enhancer function at this locus. We confirmed CTCF binding to this locus and an interaction between rs2069837 locus and a regulatory region in *GPNMB* located ∼520 kb downstream of rs2069837, suggesting CTCF-mediated long-distance chromatin looping. Indeed, we demonstrated reduced expression of GPNMB in monocyte-derived macrophages with AA genotype, which can be restored by inhibiting HDAC. A reduction in GPNMB expression in the presence of the Takayasu arteritis risk allele in rs2069837 may enhance inflammatory responses, suggesting a mechanistic pathogenic role of rs2069837 in Takayasu arteritis, while at the same time explaining a protective effect of this same allele in chronic viral infection and malignancy. Previously studies have demonstrated an anti-inflammatory role for GPNMB by enhancing M2 and suppressing M1 macrophage differentiation (*25*). The importance of the pro-inflammatory macrophages in Takayasu arteritis has been well established.

Using a combination of bioinformatics and experimental approaches, we revealed that MEF2 and HDAC proteins differentially bind to rs2069837 (more in A allele), which was highly likely to contribute to the differential regulatory function between A and G alleles. MEF2 is a transcription factor that regulates several cellular processes including differentiation, proliferation, and apoptosis (*26*). The N-terminus of MEF2 contains a highly conserved MADS-box and an adjacent motif termed as the MEF2 domain, which together mediate dimerization, DNA binding, and co-factor interactions (*27*). The co-factors binding at MEF2 N-termini determine MEF2 functions. Binding of GATA proteins will activate the expression of MEF2-target genes, while binding of HDAC IIa proteins will repress the expression of MEF2-target genes (*27*). In the present study, the same amounts of GATA proteins were observed between A allele and G allele, while multiple HDAC proteins were detected at higher levels in the presence of A allele, including HDAC IIa (HDAC4, 5, and 7). Thus the enhancer function was weakened in this locus with A allele, as we also showed using a luciferase reporter assay with and without an HDAC inhibitor. It was reported that the deacetylase activity of HDAC IIa family is not strong, but that they can recruit other robust repressors such as HDAC I proteins (*28*). Indeed, we showed significantly enriched recruitment of HDAC2 (an HDAC I family member) in the presence of the A allele in rs2069837, probably resulting from the presence of HDAC IIa which are recruited by MEF2.

The MEF2-HDAC axis plays a role in several differentiation pathways and numerous adaptive responses, including skeletal muscle differentiation, heart development, and vascular integrity (*29*). In most conditions, due to the suppression mediated by HDAC, MEF2-target genes cannot be expressed, leading to pathophysiologic alterations, as implicated in pulmonary arterial hypertension and estrogen receptor positive breast tumors (*30, 31*). Therefore, strategies to release MEF2 from HDAC proteins have been attempted. Recent findings demonstrated that inhibition of HDAC IIa or blocking the interactions of HDAC IIa and MEF2 were two potential therapeutic strategies (*32, 33*).

Existence of CTCF binding sites close to rs2069837 and CTCF enrichment at this region indicated that CTCF might mediate interactions between rs2069837 and other loci via chromatic looping. Our mass spectrometry data also showed CTCF and cohesion complex binding to this locus, further suggesting formation of chromatin loops involving this locus. Furthermore, no difference in CTCF binding between A allele and G allele was observed, implying that CTCF-mediated looping was not altered by rs2069837. Using DNase hypersensitivity maps followed by expression analysis in monocyte-derived macrophages, we identified *GPNMB* as target gene regulated by rs2069837. The 3C data further confirmed the interaction between rs2069837 locus and the regulatory region of *GPNMB*. Of interest, we did not detect a direct effect from rs2069837 on *IL6* expression, within which this polymorphism is located.

GPNMB is a negative regulator of inflammatory responses of macrophages. In dextran sulfate sodium (DSS)-induced colitis model, the deficiency of GPNMB resulted in more severe colitis characterized by higher levels of pro-inflammatory cytokines including IL-6 (*34*). GPNMB is also involved in M2 polarization of macrophages. In GPNMB-knockdown mice, M2 polarization from bone marrow-derived macrophages was inhibited, while M1 polarization and the secretion of IL-1β and TNF-α were enhanced (*35*). Therefore, it is possible that lower level of GPNMB in the presence of allele A can indirectly contribute to the expressions of multiple inflammatory cytokines including IL-6, which is important in the pathogenesis of Takayasu arteritis.

Previous genetic association studies have reported a protective effect of the A allele in rs2069837 in late-onset Alzheimer’s disease, cervical cancer, chronic hepatitis B virus infection, colorectal cancer, and hepatocellular carcinoma (*11-15*). Based on our data, the A allele in rs2069837 is associated with suppression of GPNMB expression, which is consistent with the findings that GPNMB expression is increased in Alzheimer’s disease, cancer, and infections (*36*). GPNMB weakens the immune response against cancer cells and infections, contributing to cancer development or chronic infections (*37-39*). Since GPNMB has been implicated in multiple tumors, our data discovering genetic regulation of GPNMB by rs2069837 via MEF2-HDAC complex also shed light on mechanisms that can be potentially targeted in malignant disorders.

In conclusion, our study elucidated a pathogenic mechanism underlying the genetic association between rs2069837 and Takayasu arteritis. In addition, our findings are relevant to other disorders, including chronic infection and malignancy. Targeting GPNMB and the MEF2-HDAC complex might provide novel therapeutic strategies in Takayasu arteritis. Our work highlights long-rage chromatin interactions involving regulatory variants in elucidating functional consequences of disease-associated genetic variants to pave the way for utilizing genetic data in identifying novel therapeutic approaches.

## Materials and Methods

### Cell culture

HEK293 cell line (ATCC) was cultured using Dulbecco’s Modified Eagle’s Medium (DMEM, high glucose 4.5 g/L, Hyclone) supplemented with 10% fetal bovine serum (FBS) and Penicillin-Streptomycin (100U/ml). Cells were passaged at 80-90% confluence. THP-1 cell line (ATCC) was cultured in RPMI-1640 medium supplemented with 2 mM L-glutamine, 4.5 g/L glucose, 10% FBS (not heat inactivated) at a density between 0.5 × 10^6 to 1.0 × 10^6 cells/ml. For peripheral blood mononuclear cells or monocyte-derived macrophages, RPMI-1640 supplemented with 10% FBS was used.

### Extraction and concentration detection of nuclear proteins

NE-PER™ nuclear and cytoplasmic extraction reagents (Thermo Fisher Scientific) were applied to extract nuclear protein according to the manufacturer’s manual. Next, Pierce™ BCA protein assay kit (Thermo Fisher Scientific) was used to measure the concentration of nuclear protein.

### Electrophoretic mobility shift assay (EMSA)

Thirty mer oligonucleotides flanking around rs2069837 with A/G allele (5’-TGCCAGGCACTTTA<u>A/G</u>ATAAATATTGTGTCT-3’) and their complementary oligonucleotides (5’-AGACACAATATTTAT<u>T/C</u>TAAAGTGCCTGGCA-3’) were synthesized with or without 5’ end biotin label (integrated DNA technologies [IDT], Iowa, USA). The complementary oligomers were annealed into corresponding double-stranded DNA (dsDNA) on Bio-Rad T100 Thermal Cycler (95°C for 5mins; step-cooling (95°C (−1°C /cycle), 70 cycles; holding at 4 °C). The biotin-labeled probes with A or G were incubated with nuclear extract in a binding buffer (50ng/ul Poly (dI•dC), 2.5% Glycerol, 0.05% Nonidet P-40, 5mM MgCl_2_, 10mM EDTA, 20ul system) using LightShift Chemiluminescent EMSA Kit (Thermo Fisher Scientific) for 20 min at room temperature before separating on a 6% retardation gel (Thermo Fisher Scientific). With regard to competitive EMSA, different fold excess (1-400X) of unlabeled DNA were added. After electrophoresis, the protein and DNA complexes were transferred to biodyne B nylon membrane (Thermo Fisher Scientific) and observed by chemiluminescent detection methods post-UV crosslinking.

### Prediction of binding transcriptional factors (TFs) around this locus

Different platforms were applied to analyze potential binding proteins to this locus, including HaploReg and CIS-BP database (*16, 17*). HaploReg reports ENCODE (Encyclopedia of DNA Elements) TF binding experiments (*16*). CIS-BP database collects data from >25 sources, including other databases such as Transfac, JASPAR, HOCOMOCO, and FactorBook, and predicts binding proteins based on previous experiments or similar motif across species. When 30 mer oligonucleotides used in EMSA were analyzed, several proteins were predicted to bind to the flanks and not related to the SNP of interest. Therefore, we narrowed down the input sequence to 16 oligonucleotides centering on rs2069837 with A or G (CACTTTA<u>A/G</u>ATAAATAT) to focus on the proteins most likely affected by this SNP.

### DNA affinity precipitation assay (DAPA)

Transcription factors purification was done using μMACS™ Factor Finder kit (Miltenyi Biotech, Germany). Protein-DNA binding reaction system was based on EMSA conditions and scaled up to purify enough proteins for further analysis. Specifically, around 2-2.5mg nuclear protein was incubated with 1μg labeled probe with either A, G, muted probe (5’-TGCCAGGCACTTT***GTGC***AAATATTGT GTC T-3’), or non-biotinylated oligos with A allele for 20min at room temperature. Next, 100 μl μMACS streptavidin microbeads were added to the binding reactions and incubated for 15 min at room temperature. Protein-DNA-μMACS beads complexes were then loaded onto a column in a strong magnet separator. Washing and elution processes were done following the manufacturer’s instructions. The purified proteins were identified by mass spectrometry or validated by Western blotting. In addition, about 1.5% of the input nuclear extract in this experiment was stored and later used as loading control in Western blotting.

### Identification of differential transcription factor binding by mass spectrometry

For the purified protein complex, complete reduction, alkylation, and digestion were performed by dithiothreitol (DTT, 10mM, 30min at room temperature), iodoacetamide (IAA, 10mM, dark for 30min at 37°C) and trypsin (MS Grade, Promega, 12.5ng/ul, 37°C overnight) sequentially before analyzing by Thermo Scientific™ Orbitrap Fusion™ Tribrid™ Mass Spectrometer. The proteomics data were analyzed by Proteome Discoverer software (Thermo Scientific). The number of peptide spectrum matches (PSMs) was used to compare the protein amount preliminarily. Proteins that were predicted to bind to the motif around rs2069837 and had two-fold differences between A and G were further validated by Western blotting.

### Western blotting

Proteins purified from DAPA and the corresponding 1.5% input protein (loading control) as mentioned above were added 4X Laemmli sample buffer and denatured by boiling for 10 min at 100°C on a heat block. After that, the protein samples were resolved on a 4-20% gradient SDS-PAGE and electroblotted onto nitrocellulose membranes. Various proteins selected were detected by mouse anti-GATA1, GATA2, myocyte enhancer factor 2A (MEF2A), MEF2C, CTCF, Arid3a, (histone deacetylases) HDAC2, HDAC4, HDAC5, HDAC7, and TAFII (all purchased from Santa Cruz Biotechnology) and finally developed by Super Signal West Dura substrate (Thermo Fisher Scientific) on the Omega Lum C imaging system (Gel Company).

### Luciferase reporter assays

Luciferase reporter vectors were constructed using a promoterless pRMT-Luc vector (PR100001, Origene). A 315-nucleotide fragment of the human *IL6* promoter (nucleotides–303 to +12, Ensembl ENSG00000136244) (*40*) was inserted upstream of the luciferase gene with the restriction enzyme SpeI and MluI to construct an *IL6* promoter-driven pRMT-Luc vector. In addition, a 251-nucleotide fragment flanking rs2069837 with A or G allele (hg19 chr7:22767927-22768177) were then inserted upstream of the *IL6* promoter sequence using the Xbai and EcoRI enzyme sites.

For transfection, 3×10^4^ HEK293 cells/well were seeded in a 96-well plate. Sixteen to twenty hours later, the cells were transfected with different luciferase vectors (100ng) along with 0.5ng renilla vector (pGL4.74, [hRluc/TK], Promega) using Lipofectamine 3000 reagent (Invitrogen). When THP-1 cells were transfected, SG Cell Line 4D-Nucleofector™ X Kit and Lonza Amaxa Nucleofection system were used (Program FF100) according to the manufacturer’s protocols. After 48 hours, the luminescence of firefly luciferase and renilla luciferase were detected by Synergy H1 multi-mode microplate reader (BioTek, Winooski, VT, USA) using Dual-Glo luciferase reporter assay reagents (Promega). The activity of firefly luciferase was normalized by renilla luciferase. Each condition had six replicated wells and the experiment was repeated three times. In the condition with HDAC pan-inhibitor, Trichostatin A (TSA, 400nM) or DMSO was added 2h prior to transfection and throughout the cell culture until luminescence was assessed.

### Real time PCR-based SNP genotyping

TaqMan™ SNP Genotyping Assay for rs2069837 and TaqMan™ genotyping master mix (Thermo Fisher Scientific) were used to detect rs2069837 genotype of 48 healthy subjects from whom peripheral blood mononuclear cells (PBMC) were collected and stored. 10ng genomic DNA was used for each sample, and same volume of DNase-free water was set as negative control.

### Peripheral blood monocytes isolation and differentiation into macrophages

Seven pairs of PBMC samples with AA or AG genotypes at rs2069837 were used. The age, gender, and ethnicities were matched between these two groups of subjects. Frozen PBMC were thawed quickly in a 37°C water bath and suspended in pre-warmed complete RPMI medium supplemented with 25U/ml benzonase. After washing twice, PBMC were counted and transferred to 12-well plates in 3×10^6 cells/ml with complete RPMI supplemented with 20ng/ml macrophage colony-stimulating factor (M-CSF). Two days later, the floating cells were removed, while adherent cells were washed twice and incubated with fresh medium supplemented with M-CSF for another two days. On the fourth day, fibroblast-like macrophages were obtained and media were changed again. RNA were extracted after another 24 hours. In the condition of HDAC inhibition, the media were supplemented with DMSO or HDAC pan-inhibitor TSA (100nM) for the final 24 hours.

### Extraction of messenger RNA (mRNA) and quantitative reverse transcription– polymerase chain reaction (qRT-PCR)

RNA was prepared using Direct-zol™ RNA MiniPrep kit (Zymo research, California, USA) and converted to cDNA using Verso cDNA Synthesis kit (Thermo Fisher Scientific, MA, USA). Quantitative PCR was performed with Power SYBR Green PCR Master Mix (Thermo Fisher Scientific, MA, USA) on a ViiA 7 Real-Time PCR System (Thermo Fisher Scientific, MA, USA). The primers used in the present study were listed in **Supplementary Table 1**. Gene expression data were normalized by beta-actin.

### Chromosome conformation capture (3C)

THP-1-or primary monocytes-derived macrophages were used in this experiment. 1× 10^7^ THP-1 cells were seeded in a T75 flask with complete RPMI-1640 medium (supplemented with 10%FBS) and treated with 120ng/ml phorbol 12-myristate-13-acetate (PMA) for 24 hours. After that RPMI-1640 medium without FBS and PMA was used to rest cells for 24 hours. For the primary monocytes, 10^8 PBMC were collected from healthy donors and were cultured for 4 days with M-CSF(20ng/ml) as mentioned above to obtain macrophages. For 3C experiment, we followed the protocol published by Wouter de Laat and Thierry Forné with some modifications (*41*). Briefly, per 1× 10^7^ cells, 10ml 2% formaldehyde was used to fix the cells and glycine (final concentration 0.125M) was applied to quench the formaldehyde. Then the cells were lysed to obtain, which were digested using 100U BsrDI (NEB) at 65°C overnight. 2000U T4 ligase (NEB) was applied to ligate the chromatin fragments (4h at 16°C, followed with 0.5h at room temperature) after restriction enzyme inactivation (20min at 80°C). Finally, 300ug proteinase K was used to de-crosslink the chromatin (65°C, overnight), and the ligated fragments were purified by phenol-chloroform extraction. A control template from four bacterial artificial chromosome (BAC) clones (CH17-8M1, CH17-33O24, CH17-211I22, CH17-41A22, BACPAC Resources Center) which together cover 520kb sequence spanning from *IL6* to *GPNMB* with minimally overlapping sequences were used. After culturing those clones overnight in 400ml LB Broth (Lennox, Sigma), DNA from four clones was extracted separately using nucleoBond Xtra BAC kit (Macherey Nagel). Then equimolar amounts of DNA from those clones (total 10ug) were digested using BsrDI and ligated by the T4 ligase following the conditions as mentioned above. During this process, digestion efficiency was detected by qPCR using chromatin before and after digestion with pairs of primers beside each restrictions site, and only samples that had over 65% digestion efficiency were used. In 3C-qPCR, a constant primer was designed upstream of the 5’ of BsrDI site (871bp downstream rs2069837). Paired primers for 3C detection were designed downstream of the 3’ restriction sites close to chr7:23248026 (upstream of *GPNMB*), or chr7:23288026 (in the 1st intron of *GPNMB*). The loading control was evaluated with serial dilutions of genomic DNA from THP-1-derived macrophage or the primary monocytes-derived macrophages using primers which will amplify a DNA sequence devoid of BsrDI restriction sites in *GAPDH*. The sequence of all primers used in this experiment were listed in **Supplementary Table 2**. The relative crosslinking frequencies (RCF) were normalized to both *GAPDH* loading control and BAC clones.

### Statistical analysis

Results were presented as the mean ± standard deviation (SD). The differences of luciferase signal between constructs with A and G, and gene expression in macrophages between samples with AA and AG were compared by Student’s t-test. Paired t-test was used to compare AA or AG samples with or without TSA treatment. All the analyses were performed in GraphPad Prism (Version 7.03). P-values less than 0.05 were considered significant.

### Author contributions

XK designed and performed the experiments, analyzed the data, and wrote the manuscript. AHS conceived and designed the study, designed the experiments and interpreted the data, and wrote the manuscript.

## Acknowledgements

This work was supported by the National Institute of Arthritis and Musculoskeletal and Skin Diseases of the National Institutes of Health grant number R01AR070148.

## Conflict of interest

The authors have declared that no conflict of interest exists

