## Supplementary Material for "Multi-disease associated risk locus in *IL6* represses the anti-inflammatory gene *GPNMB* through chromatin looping and recruiting MEF2-HDAC complex"

**Supplementary Figure1**

**A.** EMSA assay in THP-1 cells demonstrating increased protein binding to the probe with A allele compared to G allele. More DNA-protein complexes were shifted with the probe with allele A (lane 2) than allele G (lane 6); both unlabeled probes with A allele or G allele (24 pmol, 200X) can totally compete the corresponding shift band separately (lane 3 and lane 7); using 200X unlabeled probe with G allele can also compete the shifted band formed with the probe with A allele (lane 4); unlabeled probe with A allele was also able to compete the shifted band formed in the presence of probe with G allele (lane 8). **B.** The enhancer function of rs2069837 locus was also confirmed in THP-1 cells. Significantly increased luciferase expression was detected in both vectors with intronic sequence (A/G), with significantly higher expression in the construct with G allele than that with A allele (p< 0.05). **C.** The allele discrimination plot of 48 healthy subjects genotyped for rs2069837. **D.** No differences were observed between macrophages with AA and AG in the expressions of other genes interacting with rs2069837 based on DNase hypersensitivity correlations maps. **E.** Other epigenetic markers (H3K4me1 and H3K4me3) in monocytes also indicated the locus upstream *GPNMB* and the locus in the first intron of *GPNMB* are regulatory regions. GATA binding motif (72bp downstream of chr7:23248026 and 26bp upstream of chr7: 23288026) and MEF2 binding motif (149bp downstream of chr7:23248026 and 32bp upstream of chr7:23288026) close to these two loci are predicted, which were also demonstrated in ChIP-seq data of some cell lines (no ChIP-seq data of GATA and MEF2 in monocytes were available).

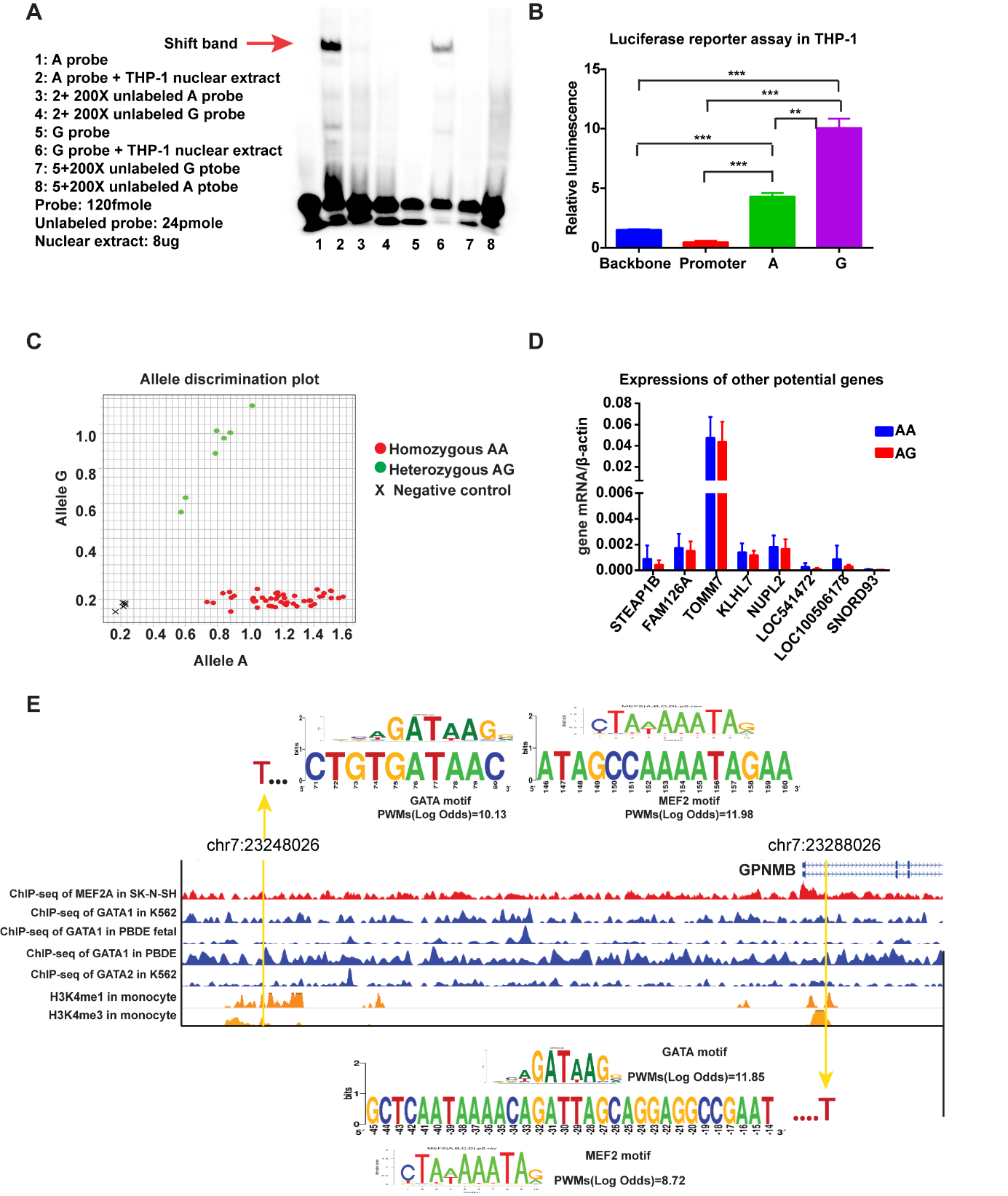

**Supplementary Table 1. Primers used in RT-PCR**

| **Gene** | **Forward primer** | **Reverse primer** |
| --- | --- | --- |
| ***Actin*** | GTCAGGCAGCTCGTAGCTCT | GCCATGTACGTTG- CTATCCA |
| ***GPNMB*** | CCTCGTGGGCTCAAATATAACAT | ACTGTCCTCTGACCATGCTGT |
| ***IL6*** | ACTCACCTCTTCAGAACGAATTG | CCATCTTTGGAAGGTTCAGG-TTG |
| ***STEAP1B*** | CCCTTCTACTGGGCACAATACA | GCATGGCAGGAATAGTATGCTTT |
| ***FAM126A*** | GTTGTGGAGGAATGGTTGTCA | GACAGGTTCTAGCAACTCACTTT |
| ***NUPL2*** | GCTTTGGATTGTCTGAGAACCC | CAAGCCTCAATTCCTCTGGTG |
| ***KLHL7*** | GTCAAGCGAGTAACACATCTTCT | AACCTGATCCTCTGCTCTCAC |
| ***TOMM7*** | CATTCGCTGGGGCTTTATCC | TCTGCACCCCTCTTAAATCCC |
| ***LOC541472*** | AAATTACTGAAGCCCACTTGGTT | ACTCTGCAAGATGCCACAAGG |
| ***LOC100506178*** | TGCAGAGCTGGTGGTCATAG | TTAGCGTGCACCAGAATGAG |
| ***SNORD93*** | ATCCTGGCCAAGGATGAGAACT | ATCCTGGCCTCAGGTAAATCCT |

**Supplementary Table 2. Primers used in 3C assay**

| **Gene** | **Primer** | **BsrDI restriction site** |
| --- | --- | --- |
| *IL6* | P_C_ : TTCTCTGATTGTCCCCCTTGA | chr7:22768898 |
|  | P_R_ : CAGCATGTTTTGGGACATTG |  |
| *GPNMB* | P1: TTCTGTGGCTGGTTTTGGT | chr7:23240228 |
|  | P2: TTCCAGAGGTGGAGGCTTTA |  |
|  | P3: GGAAAAGGTTGCTTGATTGG | chr7:23247996 |
|  | P4: CAGGCTGTAGCCCCTAGTGA |  |
|  | P5: CTGGCGGTTGGTAAACTGA | chr7:23255204 |
|  | P6: GCTACTTGGGAGGCTGAGG |  |
|  | P7: GTCAGGCTGGTCTCGAACTG | chr7:23254931 |
|  | P8: AGAGCCTTTTACACCATTCCA |  |
|  | P9: TGGTGGTGTGTGCCTGTAGT | ch7:23282317 |
|  | P10: TTCATGTAATGTAAAGCGAATAATG |  |
|  | P11: CAGGAGAATGGCTTGAACCT | chr7:23286092 |
|  | P12: GACTCACTCCTTCTTGGCATCT |  |
|  | P13: GGTGGTGTGTGCCTGTAGT | chr7: 23288603 |
|  | P14: TTCATGTAATGTAAAGCGAATAATG |  |
|  | P15: GGTGAAGTCCCGTCCCTACT | chr7: 23297260 |
|  | P16: GCTCATCAATGTGCCATTCT |  |
| *GAPDH** | F: ACAGTCCATGCCATCACTGCC |  |
|  | R: GCCTGCTTCACCACCTTCTTG |  |

Pc: constant primer; * primers in *GAPDH* were used in loading control assessment. Primers pairs next to a restriction sites were used to evaluate digestion efficiency (Pc and P_R_, P1 and P2, P3 and P4, P5 and P6, P7 and P8, P9 and P10, P11 and P12, P13 and P14, P15 and P16), while the constant primer paired with P2, P4, and P6, P8, P10, P12, P14, P16 respectively were used in 3C analysis.

**Supplementary Table 3. Regulatory motifs altered on rs2069837**

| **Position Weight Matrix ID** (Library from [Kheradpour and Kellis, 2013](http://compbio.mit.edu/encode-motifs/)) | **Strand** | **Ref** | **Alt** | **Match on:**  **Ref:** TAAGTATCTACTGTGTGCCAGGCACTTTAAATAAATATTGTGTCTAATCTTCAAAACAA **Alt:** TAAGTATCTACTGTGTGCCAGGCACTTTAGATAAATATTGTGTCTAATCTTCAAAACAA |
| --- | --- | --- | --- | --- |
| **Arid3a_2** | + | 11.6 | 6.1 | NNNNTTRATYAAWHNHN |
| **Arid5a** | - | 11.5 | 10.5 | DNYHBHAATATTRB |
| **Fox** | - | 16 | 12.4 | WAARYAAAYAWWV |
| **Foxa_known1** | - | 12.9 | 10.2 | HWRWRYAAAYA |
| **Foxa_known4** | - | 14.7 | 10.6 | WRARYAAAYAWKNMV |
| **Foxd3** | - | 12.8 | 8.5 | RAAHMAAYAWWY |
| **Foxf1** | - | 14.1 | 10.1 | YRHAYAAACAHNB |
| **Foxi1** | - | 11.6 | 7.4 | WAWRYAAAYAHVH |
| **Foxj1_1** | - | 11.9 | -0 | TAAACAAACAHWD |
| **Foxj2_1** | - | 14.5 | 12.7 | HHNHMRRYAAAYAHHNNW |
| **Foxl1_1** | - | 14.8 | 13.8 | DHDVHATAAAYAHDDN |
| **HDAC2_disc2** | + | 11.9 | 10.4 | WRRGYMAAYA |
| **Hlx1** | - | 11.4 | 8.1 | DRTAWTYAAWTADKD |
| **Mef2_known5** | + | 6.9 | -5 | HSTGTTRCTAWAAATAGAWHMN |
| **Nkx6-2** | - | 11.8 | 10.8 | RWRDTAAWTABB |
| **TATA_known1** | + | 12 | 9.1 | NHDWWWTWWAWWWDRN |
| **TATA_known3** | + | 10.8 | 11.3 | KATAAATW |

Ref: reference allele; Alt: alternative allele; values in the table were log-odds scores of position weight matrices (PWMs). S = C or G, W = A or T, R = A or G, Y = C or T, K = G or T, M = A or C, N = any base pair.

**Supplementary Table 4. Predicted binding proteins in the rs2069837 locus using CIS-BP database**

| Name | Motif ID | Family | Sequence | From | To | Direction | Score-A | Score-G |
| --- | --- | --- | --- | --- | --- | --- | --- | --- |
| ARID3 | M0106_1.02 | ARID/BRIGHT | AAATAAATA | 7 | 15 | R | 8.04 | - |
| FOXA | M6241_1.02 | Forkhead | TAAATAAATA | 6 | 15 | F | 14.932 | 13.313 |
| FOXD | M6241_1.02 | Forkhead | TAAATAAATA | 6 | 15 | F | 14.932 | 13.313 |
| FOXI | M6241_1.02 | Forkhead | TAAATAAATA | 6 | 15 | F | 14.932 | 13.313 |
| FOXJ | M6241_1.02 | Forkhead | TAAATAAATA | 6 | 15 | F | 14.932 | 13.313 |
| FOXL | M6241_1.02 | Forkhead | TAAATAAATA | 6 | 15 | F | 14.932 | 13.313 |
| MEF2 | M6340_1.02 | MADS box | ACTTTAAATAAATA | 2 | 15 | F | 10.525 | - |
| GATA | M3314_1.02 | GATA | TTTAGATAAATAT | 4 | 16 | F | - | 8.627 |

F**:** forward; “From” and “to” mean the position of the sequence CACTTTAA/GATAAATAT which matched the corresponding motif; The “score” evaluates each position in the sequence with all PWMs, using a standard log odds scoring method (Nat Biotechnol. 2011;29(6):480-3.).

**Supplementary Table 5. Binding proteins observed with mass spectrometry. The list is filtered to include only proteins detected using data included in Supplementary Tables 2 and 3 (See Methods).**

| **Family/protein** | **Protein name** | **Coverage** | **Unique Peptides** | **MW [kDa]** | **PSMs (unlabeled)** | **PSMs (Muted)** | **PSMs (A)** | **PSMs (G)** |
| --- | --- | --- | --- | --- | --- | --- | --- | --- |
| **ARID** | Arid3A | 25.29 | 6 | 62.9 | 1 | 1 | 7 | 5 |
|  | Arid3B | 19.25 | 8 | 60.6 | 1 | 3 | 9 | 5 |
| **Fox** | FOXJ3 | 20.25 | 7 | 68.9 | 2 | 6 | 10 | 5 |
|  | FOXA1 | 46.39 | 9 | 49.1 | 0 | 8 | 16 | 12 |
| **MEF2** | MEF2A | 7.49 | 1 | 54.8 | 0 | 0 | **3** | **0** |
|  | MEF2C | 7.61 | 1 | 51.2 | 0 | 0 | **3** | **0** |
|  | MEF2D | 8.06 | 2 | 55.9 | 0 | 0 | **4** | **0** |
| **GATA** | GATA1 | 26.92 | 6 | 42.7 | 1 | 4 | **2** | **8** |
|  | GATA2 | 27.39 | 8 | 50 | 2 | 3 | **2** | **12** |
| **TAF** | TAF1 | 4.64 | 7 | 212.5 | 1 | 5 | 1 | 3 |
|  | TAF4 | 14.74 | 8 | 110 | 2 | 7 | 3 | 4 |
| **HDAC** | HDAC2 | 56.35 | 14 | 55.3 | 2 | 19 | 19 | 15 |
|  | HDAC5 | 25.70 | 6 | 121.9 | 0 | 4 | **3** | **0** |
|  | HDAC7 | 61.61 | 14 | 102.9 | 4 | 20 | 19 | 20 |

PSMs: peptide spectrum matches; MW: molecular weight; ARID: AT-rich interaction domain containing (ARID) family; Fox: forkhead box family; MEF2: myocyte enhancer factor-2; TAF: TATA box binding protein associated factor; HDAC: Histone deacetylase.

**Supplementary Table 6. Binding proteins detected by mass spectrometry that also participate in chromatin looping**

| **Family/protein** | **Protein name** | **Coverage** | **Unique Peptides** | **MW [kDa]** | **PSMs (unlabeled)** | **PSMs (Muted)** | **PSMs (A)** | **PSMs (G)** |
| --- | --- | --- | --- | --- | --- | --- | --- | --- |
| **CTCF** | CTCF | 41.67 | 22 | 82.7 | 3 | 16 | 12 | 11 |
| **Cohesin**  **complex** | SMC1A | 62.53 | 81 | 143.1 | 2 | 66 | 52 | 66 |
|  | SMC3 | 61.54 | 79 | 141.5 | 4 | 63 | 61 | 64 |
|  | RAD21 | 63.07 | 30 | 71.6 | 2 | 20 | 15 | 26 |
|  | STAG1 | 14.78 | 10 | 144.3 | 3 | 7 | 8 | 6 |
|  | STAG2 | 39.88 | 30 | 141.2 | 5 | 23 | 25 | 19 |

PSMs: peptide spectrum matches; MW: molecular weight; CTCF: CCCTC-binding factor.

**Supplementary Table 7. Annotation analysis of the genes interacting with rs2069837 using DNase hypersensitivity data**

| **Category** | **Term** | **Count** | **%** | **P-Value** | **Genes** |
| --- | --- | --- | --- | --- | --- |
| GOTERM_BP_DIRECT | GO:0045765~regulation of angiogenesis | 2 | 16.6 | 0.015 | *IL6, GPNMB* |
| GOTERM_BP_DIRECT | GO:1901215~negative regulation of neuron death | 2 | 16.6 | 0.018 | *IL6, GPNMB* |
| GOTERM_BP_DIRECT | GO:0070374~positive regulation of ERK1 and ERK2 cascade | 2 | 16.6 | 0.080 | *IL6, GPNMB* |
| GOTERM_MF_DIRECT | GO:0005515~protein binding | 8 | 66.6 | 0.045 | *KLHL7, IL6, TOMM7, MALSU1, IGF2BP3, NUPL2, GPNMB, FAM126A* |

**Supplementary Table 8. Demographic information of the healthy subjects using in this study**

| **Sample** | **Gender** | **Ethnicity** | **Age** | **Genotype of rs2069837** |
| --- | --- | --- | --- | --- |
| 1 | Female | Hispanic | 22 | AA |
| 3 | Female | Caucasian | 22 | AA |
| 4 | Male | Caucasian | 23 | AA |
| 5 | Female | Caucasian | 25 | AA |
| 6 | Female | Caucasian | 52 | AA |
| 7 | Female | Caucasian | 53 | AA |
| 2 | Female | Caucasian | 64 | AA |
| 8 | Female | Hispanic | 20 | AG |
| 9 | Female | Caucasian | 22 | AG |
| 10 | Male | N/A | 22 | AG |
| 11 | Female | Caucasian | 25 | AG |
| 12 | Female | Caucasian | 54 | AG |
| 13 | Female | Caucasian | 56 | AG |
| 14 | Female | Caucasian | 64 | AG |
